# Star-polymers as potent broad-spectrum extracellular virucidal antivirals

**DOI:** 10.1101/2024.07.10.602907

**Authors:** Elana H. Super, Si Min Lai, Urszula Cytlak-Chaudhuri, Francesco Coppola, Olivia Saouaf, Ye Eun Song, Kerriann M. Casey, Lauren J. Batt, Shannan-Leigh Macleod, Robert H.T. Bagley, Zarah Walsh-Korb, Petr Král, Eric A. Appel, Mark A. Travis, Samuel T. Jones

## Abstract

Viruses pose a significant threat to both global health and the global economy. It is clear that novel antiviral strategies are urgently needed, with a broad-spectrum approach being most desired. We have discovered a broad-spectrum, non-toxic polymer virucide that can tackle the viral threat. This polymeric virucide is effective at nanomolar concentrations, against a broad-spectrum of viruses and, demonstrated using an intranasal respiratory syncytial virus (RSV) murine model, has excellent efficacy, low anti-coagulant properties and low toxicity *in vivo*. Molecular dynamic simulations show that this polymer achieves its virucidal antiviral effect *via* self-assembly of viral-receptors leading to increased envelope forces and viral disassembly. The discovery of this cheap and readily produced polymer marks the start of a new type of receptor-crosslinking broad-spectrum antiviral that has significant potential to combat the global threat posed by viruses.

---

Viruses are a potential threat to every known species on the planet. These obligate parasites cause a wide range of infections, mutate rapidly and are widely untreatable. Respiratory viruses, in particular, have historically had some of the greatest impacts on human life either through direct infection or infection of animal/food stocks.^1–4^ They are capable of human-to-human as well as zoonotic transmission, allowing them to rapidly spread across the globe.^1–8^ Influenza and coronaviruses, two well-known respiratory viruses, have recently had devastating effects on both human^1, 2, 9^ and animal populations.^5, 6^ Containing viruses, especially respiratory viruses, is of critical importance to public health, the global economy, and societal growth and maintenance. It is also imperative that any interventions are inexpensive and easily distributed to lower to middle income countries, which can act as viral sinks during outbreaks.^10–14^

Although vaccination currently prevents some viral infections, therapeutics are of great importance, especially when there is no vaccine available^12–18^ or for those already infected. Typically, small molecule drugs that target a specific stage of the viral replication cycle are developed as antiviral therapeutics. These are virus-specific drugs to which viruses can become resistant, on account of rapid viral mutations. Where no effective antivirals are available, monoclonal antibodies (MAbs) may also be a potential therapeutic route. Antibodies are often virus-specific but have long circulation life-times and strong viral binding.^19, 20^ However, their high cost^21–24^ and the lack of pan-viral antibodies^21–23^ limits their use and accessibility.

For wide-reaching protection against viruses, an antiviral with broad-spectrum efficacy would be most effective however, no approved broad-spectrum antiviral treatment options exist.^25^ However, there are many well-known substances with broad-spectrum antiviral properties. Disinfectants, such as bleach, oxidising agents and alcohols, are broadly acting virucides (destroy viruses on contact) (Figure 1A). In recent years, the effectiveness of such virucides at controlling viral spread has been widely proven *i*.*e*. hand sanitisers. While they are effective at destroying viruses, they are also highly toxic and potentially flammable. Soaps/surfactants also have well-documented virucidal properties,^26, 27^ with recent work on surfactant-coated cores (nanoparticle, dendrimers, cyclodextrins and small molecules) also showing promise as broad-spectrum virucides (Figure 1B).^25, 28–30^ However, surfactants are also toxic, as well as damaging to the environment, limiting their broad application. If a non-toxic broad-spectrum virucide were to exist, it could have significant potential for the prevention and treatment of many viral infections. Polymers have significant potential advantages for the treatment and prevention of viruses, including ease of synthesis, low cost, and extended shelf-lives/stability. Both natural and synthetic sulfonated polymers are known to interact with viruses and are generally non-toxic.^31–35^ However, the study of polymers as antivirals, and their virustatic (reversible inhibition) antiviral mechanism, is well-documented and prohibits their use clinically.^36–41^ Many also inhibit blood coagulation (i.e heparin),^42, 43^ making this a key consideration for any developed sulfonated polymer antivirals.

**Figure 1.**
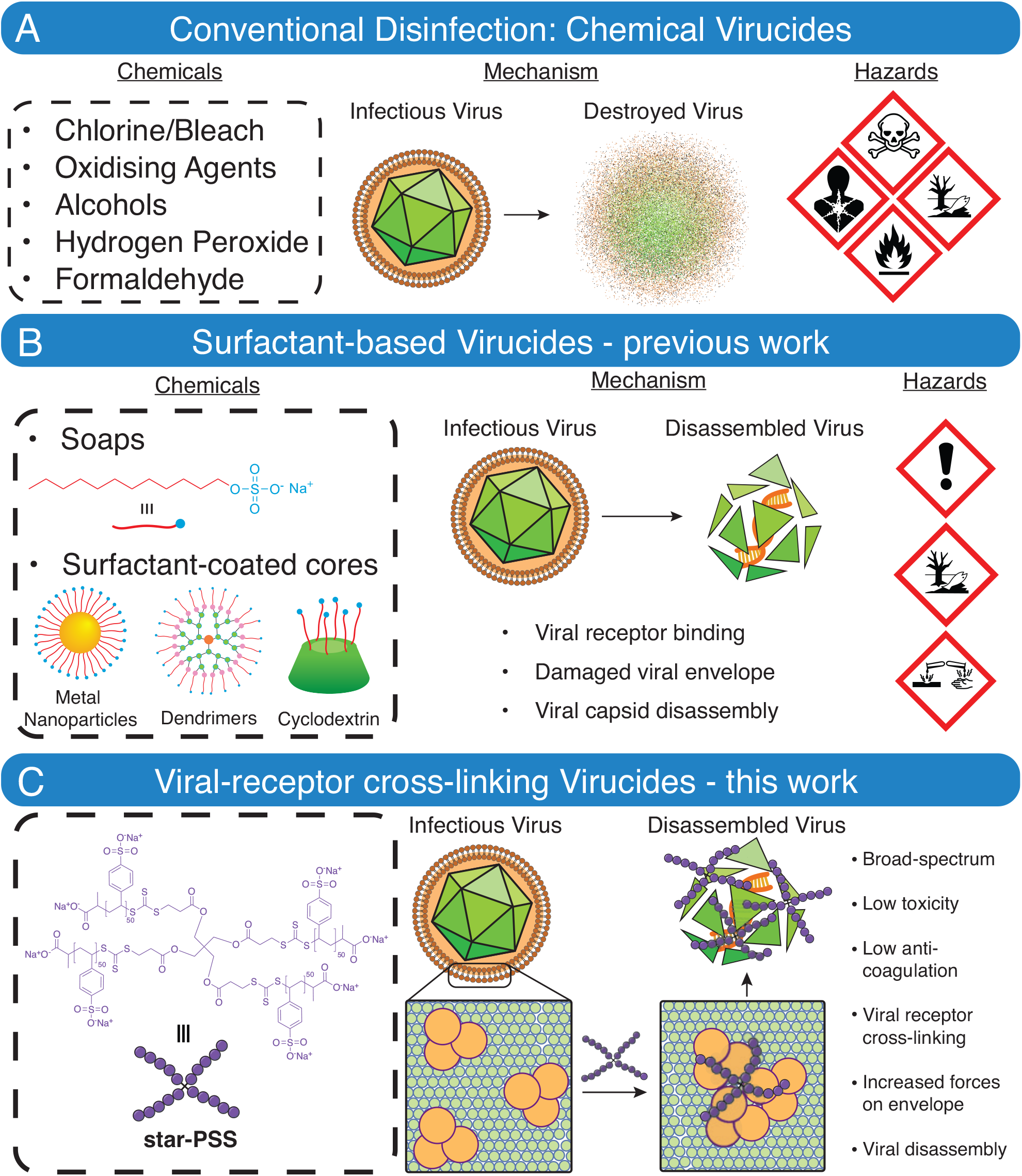
Disinfectants are powerful broad-spectrum antivirals but are typically toxic. New biocompatible virucides are needed. A) Example of conventional chemical virucides that destroy all components of viruses through chemical reactions (i.e. oxidation), but are known to be highly toxic, B) Surfactants and surfactant-based virucides cause viral disassembly by interrupting intermolecule interactions (i.e. hydrophobic interactions between lipids) but known to have toxic properties and C) Novel viral-receptor cross-linking approach that increases forces applied to envelopes, leading to viral disassembly, without observed toxicity.

We set out to explore why such a promising class of antiviral has failed to achieve the broad-spectrum and highly-desired virucidal (destroy-on-contact) mechanism. We hypothesised that the structure-property relationship between linear sulfonated polymers and viral binding proteins did not allow for sufficient interactions to induce a virucidal effect. We hypothesised that changing the architecture of these sulfonated polymers, from linear to multi-arm star, could increase the chance of sufficient interactions between the virus and polymer, thereby enhancing the antiviral mechanism.

In order to achieve a star polymer architecture, aqueous reversible addition-fragmentation chain transfer (RAFT) polymerisation, using a carboxylic acid-terminated multi-arm chain-transfer agent (CTA) with 4-arms (one CTA per arm), was used.^44, 45^ When utilised alongside sodium 4-vinylbenzene sulfonate, sulfonated 4-arm star polymers of poly(styrene sulfonate) (PSS) are produced, termed **star-PSS**, from one 4-arm CTA (Figure 1C). On account of our previous work,^46, 47^ and that of others,^35, 48–50^ that explore PSS as an antiviral, we synthesised polymers with a degree of polymerisation (DP) per arm of 50. This size of polymer has previously been shown to have low half-maximal effective concentration (EC_50_) values against a wide array of viruses with very low toxicities.^32, 46^ DP50 **star-PSS** was synthesised *via* a simple overnight reaction in water, dialysed and lyopholised before being fully characterised (Figure S1 and S2). The synthesised polymers show a relatively broad PDI for a RAFT polymerisation (Figure S1), and could not be further reduced, which is likely a result of the star architecture.^44, 51^ Nevertheless, these polymers were simple to produce using an environmentally friendly solvent (water), in high yield. DP50 **star-PSS** was used in all further studies.

Initial evaluation of the **star-PSS** was focused on determining if the broad-spectrum antiviral and low toxic properties of linear PSS (**L-PSS**), a known virustatic polymer, had been maintained.^36, 38, 40, 41, 46^ MTT assays (Figure S3) were used to determine the toxicity of **star-PSS**, which clearly showed no toxic effects, even up to 5 mg/mL of direct contact between polymer and cells. Determining the cell toxicity is a critical first step in antiviral assessment, as all further steps require the use of cultured cell monolayers and any cell toxicity would exclude materials from further testing.

Dose response assays, utilising plaque counting, were used to determine EC_50_ values against a range of viruses. Figure 2A shows the EC_50_ values for **star-PSS** against 5 different viruses - respiratory syncitial virus (RSV), Herpes Simplex Virus-2 (HSV-2), Human Coronavirus strain OC43 (OC43), Human Rhinovirus 8 (HRV8) and Cytomegalovirus (CMV). It can clearly be seen that **star-PSS** is a highly efficacious broad-spectrum antiviral with EC_50_ values typically in the nano-to pico-Molar range. The affect of **star-PSS** against CMV is considerably worse (higher EC_50_), but still efficacious at low concentrations, the reason for this difference is currently under investigation but may be linked to the mechanism of action (*vide infra*). Selectivity indices (SI) are often used to compare antivirals, this is the ratio between the half-maximal cytotoxicity concentrations (CC_50_) and EC_50_ values. On account of the extremely low toxicity and antiviral efficacies, **star-PSS** has some of the highest SI values ever reported, *e*.*g*. SI value against RSV of 325,662. This is orders of magnitude greater than any previously reported antiviral against RSV. It should be noted that the calculated SI values in Figure 2A are calculated with a CC_50_ value of 300 *µ*g/mL, which we have selected as the upper limit of toxicity for this study. As can be seen in Figure S3, no toxicity was observed for **star-PSS** even up to 5 mg/mL, and likely could be further increased. This means that the SI values could be recalculated with this highest tested concentration, greatly increasing the SI values to unprecedented levels, *e*.*g*. RSV = 5,427,703.

**Figure 2.**
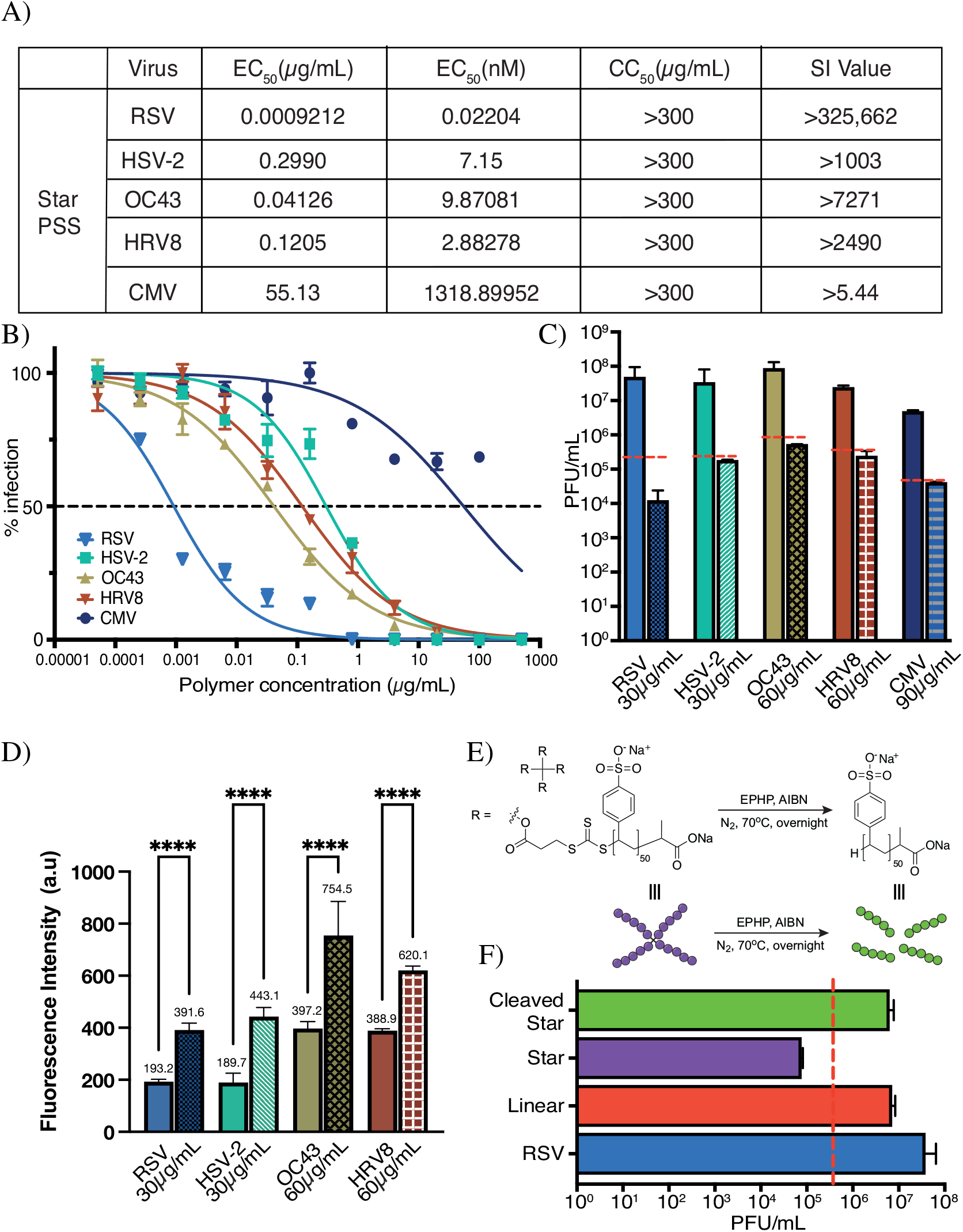
star-PSS is a highly effective broad-spectrum antiviral with a virucidal mode of action. Data overview for the testing of **star-PSS** against respiratory syncytial virus (RSV), herpes simplex virus 2 (HSV-2), human coronavirus OC43 (OC43), human rhinovirus 8 (HRV-8) and cytomegalovirus (CMV), showing A) the highly efficacious, broad-spectrum effect with low EC_50_’s and high selectivity index (SI) values, B) dose response assay data, C) shows that **star-PSS** is a broad-spectrum virucide with left bar being no treatment control (NTC) and right treated (red line = 2-log reduction, indicating a virucidal material), D) fluorescent FAIRY assay^52^ results showing statistically significant increases in fluorescence, before (left) and after (right) mixing with **star-PSS**, confirming a virucidal mechanism, and E) schematic of the cleavage of **star-PSS** to yield **L-PSS** alongside (F) viral plaque counting data for **L-PSS** vs **star-PSS**, confirming the star architecture is necessary for a virucidal mechanism. Significance was calculated using a One Way ANOVA with multiple comparisons, n = 3. **** = p<0.0001.

Once the low toxicity and antiviral efficacy of the **star-PSS** had been confirmed, it was necessary to investigate the antiviral mode of action. For this, a plaque-based dilution study was used following a well-known and reported virucidal assay.^25, 29, 46, 47, 53^ Here we attempt to induce a virustatic response to an antiviral through a heavy dilution of the virus-antiviral interaction. The assay indicates a virucidal effect when a 2-log reduction in virus is detected, when compared to a no treatment control (NTC). It can be seen in Figure 2C that **star-PSS** has the desired virucidal mode of action (*>*2-log decrease in PFU/mL - red dotted line) against each of the viruses tested. To further confirm the virucidal mode of action, a FAIRY assay,^52^ which assesses fluorescence increases upon release of viral genomes (virucidal), was used. Statistically significant increases in fluorescence were obtained for all viruses tested, further confirming the virucidal mechanism (Figure 2D).

To confirm that it was the polymer architecture that was responsible for the change in mode of action, the **star-PSS** was cleaved back to linear PSS. The trithiocarbonate, embedded within each arm of the star polymer, can be readily cleaved. As this group remains at the core of the star structure, the resultant polymers, after cleavage, are linear PSS (Figure 2E). It can be seen in Figure 2F that following cleavage of the core (producing linear PSS) the antiviral mechanism returns to a virustatic one. This confirms that the star shape is key to the virucidal mode of action.

The virucidal nature of **star-PSS** is highly appealing, however, its viral deactivating mechanism was unclear. We hypothesize that **star-PSS** acts like a molecular glue, which has recently been proposed,^54^ assembling the protein receptors within the viral membrane,^55, 56^ similar to the effect of negative lipid membrane curvature during virus endocytosis.^57^ To test this hypothesis, we performed molecular dynamics (MD) simulations of **star-PSS** (total 200 sulfonates) and linear-50 and 200 **L-PSS** polymers coupled to individual and grouped viral receptors (See Methods). First, the polymers were simulated in a physiological solution in contact with one viral (RSV) receptor protein (Figure S4). RSV proteins were used during *in silico* experiments as they are well defined and **star-PSS** showed the highest efficacy against RSV *in vitro*.

The **star-PSS** perfectly wrapped the individual proteins, and provided a large binding energy of *-*296.5 kcal/mol, while the linear-50 and 200 PSS had smaller binding energies of *-*116.1 kcal/mol and *-*169.1 kcal/mol, respectively. Although, these results revealed a higher activity in **star-PSS** (see video S1), they didn’t explain their virucidal behavior.

To examine if the hypothetical protein assembly is a realistic possibility, we simulated two polymers of each type coupled to a loose aggregate of 7 RSV receptor proteins placed in a fictive membrane. These 7 proteins were attached by their bottom atoms to a planar constriction allowing their lateral sliding along the ‘membrane’ (Method). Figure 3 indeed reveals that the receptor proteins become aggregated when either the star or linear polymers are present. However, the activities of these two polymers are significantly different. In particular, the **star-PSS** polymers behave like a molecular glue, each capable of attracting 3 proteins, as seen in Figure 3A-C. In contrast, **L-PSS** (DP = 200) polymer can only couple pairs of proteins, as seen in Figure 3D-F. Figure 3G,H also provides the average forces that these polymers exert on the proteins. They are obtained as fits to distance-dependent binding energies between the polymers and proteins (Methods). This analysis shows that the **star-PSS** polymer exerts about a twice larger force than the **L-PSS** (DP = 200) polymer (*F ≈* 850 pN vs *F ≈* 480 pN), it can not only assemble the proteins (video S2), but it can also deform the underlying lipid membrane, thus, compromising its stability. In this way, **star-PSS** polymers could be virucidal. We have shown that **star-PSS** has very low toxicity *in vitro* (Figure S3) with a virucidal mode of action (Figure 2C). Any potential treatment option for respiratory viruses, that relies on direct contact with a virus (*i*.*e*. virucidal), would need to be administered intranasally. In order to confirm that **star-PSS** maintains its low *in vitro* toxicity profiles *in vivo*, intranasal toxicity assays were used. Mice repeatedly treated with **star-PSS** generally displayed minimal, insignificant toxicity compared to control subjects. Within serum biochemistry panels, we found that alanine transaminase (ALT), aspartate transaminase (AST), bilirubin, and creatinine were within expected ranges and did not differ between control and treatment groups. Blood urea nitrogen (BUN) was favourably lowered in antiviral-treated mice Figure 4A). Similarly, histopathology indicated no systemic lesions in the heart, liver, or kidneys but there was minor damage at the local site of inoculation, the nasal turbinates (Figure 4B - black arrow). The changes were mild and scattered throughout the caudal most section (section 4) of the nasal turbinates within each mouse. Histologic changes included erosive and neutrophilic rhinitis and variable apoptosis of the respiratory and olfactory epithelium. In rostral sections (sections 1-3), treated mice displayed mild symptoms of increased mucus production and scattered cellular debris within the lumen, likely a result of damage within the caudal turbinates. Overall, no adverse findings were observed following intranasal administration of **star-PSS**, further confirming the potential for treatment of respiratory viruses, such as RSV.

**Figure 3.**
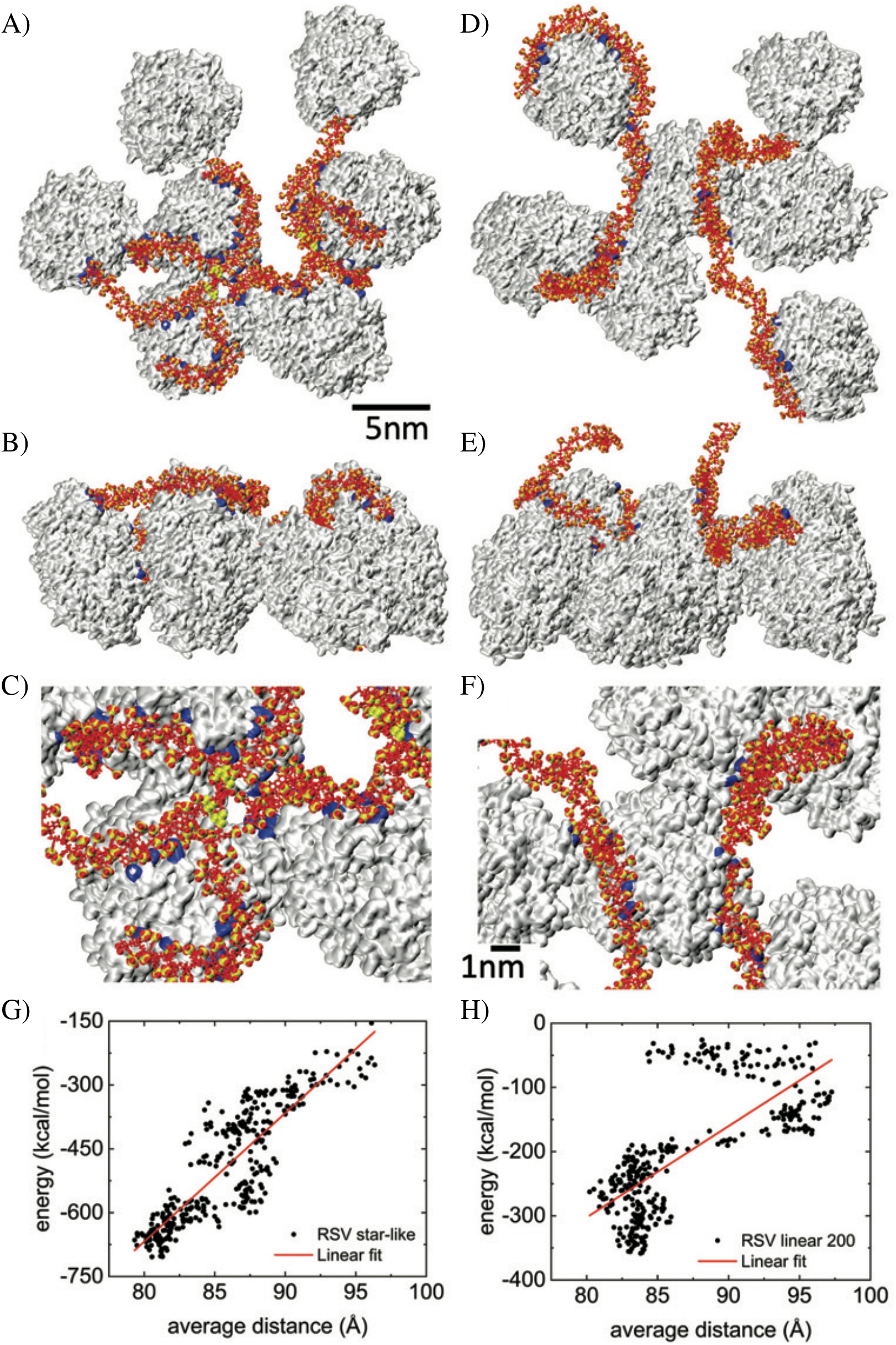
star-PSS crosslinks viral-surface receptors leading to increased viral membrane forces. Molecular dynamic simulations showing a pair of (A-C) **star-PSS** (DP = 50) and (D-F) **L-PSS** (DP = 200) polymer coupled to 7 RSV receptor proteins for 350 ns of simulations (C, F details). In blue, basic residues within 4 Å from the polymers. (G, H) The protein-distance dependence of the coupling energies provide the forces with which the polymers act on the proteins and assemble them. In red, the fit lines used to extract the force from the slope.

**Figure 4.**
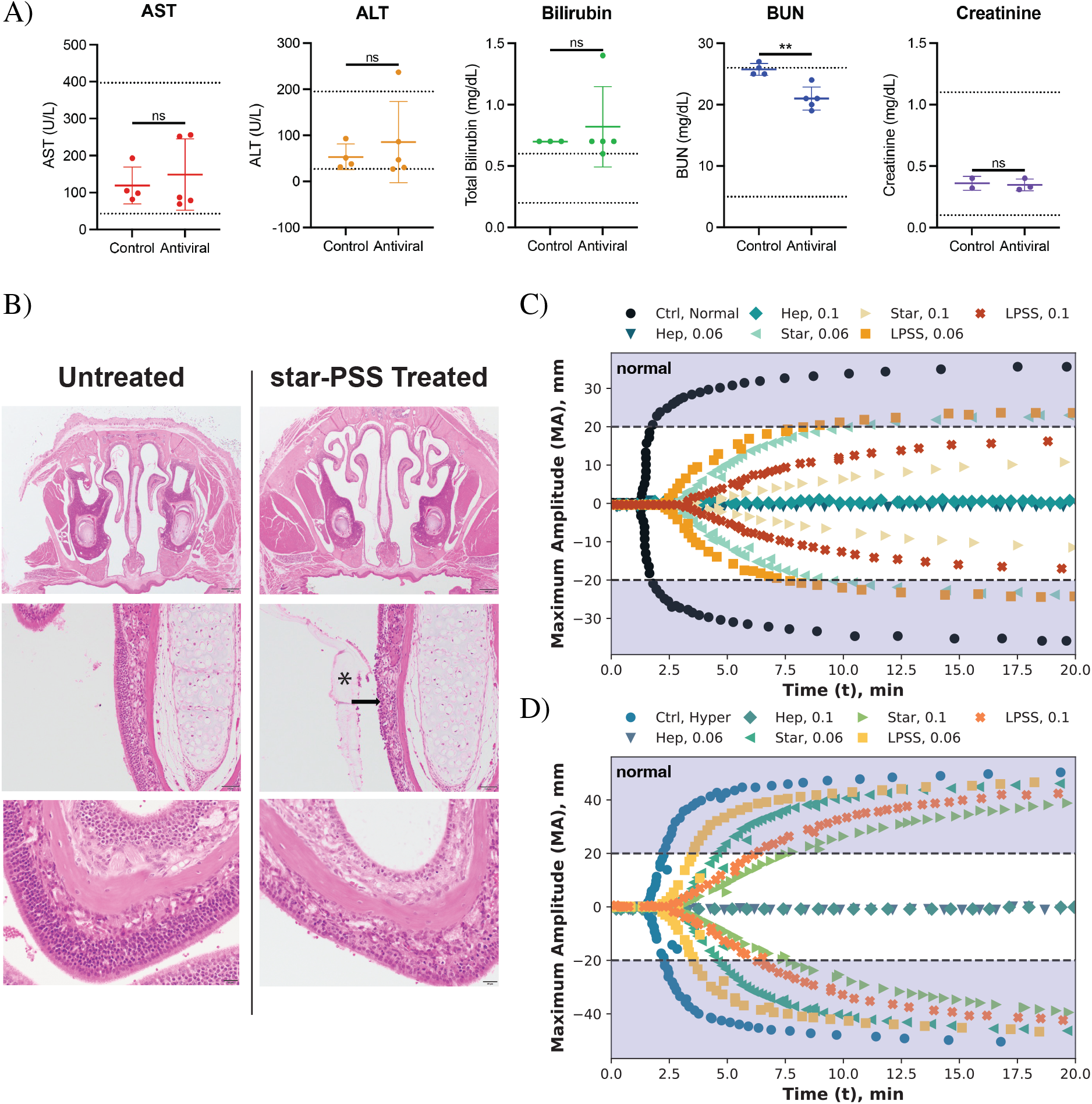
star-PSS shows no blood panel toxicity and minimal damage at the site of administration following intranasal *in vivo* administration and it is not an effective anti-coagulant. A) Blood panels for alanine transami-nase (ALT), aspartate transaminase (AST), bilirubin, blood urea nitrogen (BUN), and creatinine following 100*µ*g **star-PSS** for 5 days in Mice (n = 5, C5BL/6, female), B) histopathology of mouse turbinates demonstrating mild erosive and neutrophilic rhinitis (arrow) with luminal mucus and neutrophilic debris (asterisk) in **star-PSS** treated mice, thromboelastography study into the anti-coagulation properties of **L-PSS** and **star-PSS** vs heparin sulfate (at 0.06 mg/mL and 0.1 mg/mL) in both normal (C) and hyper (D) coagulating blood.

On account of the similarity to heparin (a known anticoagulant), sulfonated star polymers may exhibit anti-coagulant properties. We used thromboelastography (TEG) to evaluate how **star-PSS**, as well as **L-PSS**, affects blood coagulation, focusing on clotting onset time (R, min), the time to reach the minimum viable clot strength (K, min), the maximum amplitude or maximum clot strength (MA, mm), and clot formation rate (*a*, degrees).^58–60^ Both normal and hypercoagulable (HC) blood samples were tested with **star-PSS** and **L-PSS** at 0.06 mg/ml (equivalent to heparin conc. in blood collection tubes^61^) and 0.1 mg/ml (concentration used in *in vivo* toxicity studies) and the results are shown in Figure 4C and D. Hypercoagulability, where blood tends towards heavy clotting, is particularly common after injury or infection, affecting approximately 1 in 500 people.^62–64^

In normal blood, 0.06 mg/ml **star-PSS** slightly prolonged R and K and reduced *a* and MA, showing minor anti-coagulation effects without preventing viable clot formation (Figure 4C, turquoise triangles). At 0.1 mg/ml, clot strength was insufficient, although the typical clotting pattern was maintained (Figure 4C, yellow triangles). This suggests that the antivirals minimally impacts clot formation by interfering with the ability for the clotting factors to bind to their targets, rather than in any other way (e.g., precipitation of clotting factors). In HC blood, 0.06 mg/ml **star-PSS** or **L-PSS** had minimal impact, with the MA still well above viable levels. At 0.1 mg/ml, effects were more pronounced but did not critically impair clot strength (Figure 4D).

HS is a linear polymer, while **star-PSS** is a 4-arm star. We therefore wanted to understand if there is a structural reason for the lower impact of **star-PSS** on clotting compared to linear HS or PSS. As discussed above, **star-PSS** can be cleaved to a linear PSS equivalent. From Figure 4C, where the maximum values for each parameter are given, the linear PSS appears to have less of an impact on the four clotting parameters than the **star-PSS** (Figure 4C, red and orange squares). This was unexpected as it was hypothesised that the linear PSS would behave more similarly to the linear HS and have a greater negative impact on clotting behaviour. The cause for this deviation is likely related to the lower affinity of the **L-PSS** for the heparin binding sites on the blood coagulation factors, in particular thrombin, which we observe in MD simulations (Figure S5-7). Overall, this data clearly suggests that **star-PSS** is an effective antiviral but is not an effective anti-coagulant.

Given the potent antiviral activity of **star-PSS** *in vitro*, we wanted to test the efficacy of the polymer during viral infection *in vivo*. To this end, we utilised a murine model of RSV infection, one of the most prevalent and detrimental respiratory viruses for which no antiviral treatment options are available.^11, 65^ RSV, through annual outbreaks, accounts for an estimated 33 million severe infections per year, including around 3 million hospitalisations and 120,000 deaths.^66–73^ To add to the public health impact of RSV, over 95% of the occurrences of severe RSV-induced pneumonia and bronchiolitis effect lower to middle income countries.^11, 73^ Of the 120,000 annual mortalities, over 99% of these are found in these same lower to middle income countries,^11, 72–74^ meaning that any intervention developed against RSV, while critical, must be readily available to those in lower to middle income countries.

Mice were infected with RSV by the intranasal (i.n.) route, and from 24 hours post-infection were treated with 25 *µ*g **star-PSS** (or PBS as control) intranasally daily for 3 days (Figure 5A). We found that treatment with **star-PSS** caused a profound reduction in lung viral titres compared to PBS-treated mice, with a nearly 4-log reduction, which equates to a viral load reduction of 99.99% (Figure 5B). To determine whether polymer treatment had any effect on the host immune system during infection, we analysed pulmonary damage (Figure 5C) and lung immune cells by flow cytometry (Figure 5 D-I and Figure S10 B-H). Treatment with **star-PSS** during infection did not alter total CD45+ immune cell numbers in the lung (Figure 5D), indicating that the polymer was not causing any alteration in the immune response to infection. In support of this, we did not observe any alteration in numbers of alveolar macrophages or monocytes (Figure 5D-E), nor the cDC1 dendritic cell subset, TCR*b* + T cells, *gd* T cells, B cells, eosinophils, or NK cells by **star-PSS** treatment in the lung during RSV infection (Figure S10B-H). We did observe a significant increase in inflammatory macrophage, cDC2 dendritic cell, and neutrophil numbers induced by **star-PSS** treatment during infection (Figure 5G-I).

**Figure 5.**
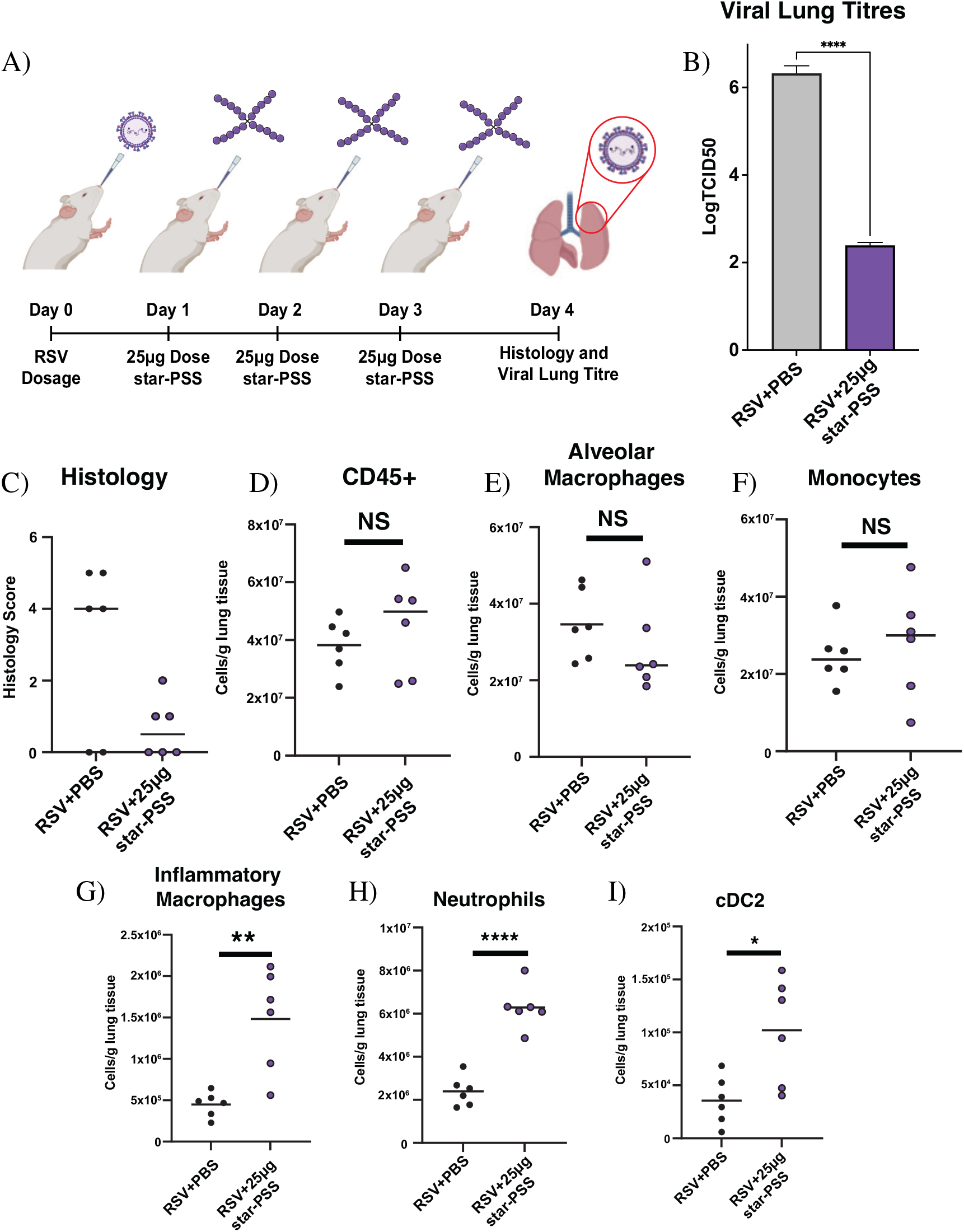
In vivo treatment with polymer reduces viral load and inflammation during RSV infection. A) Schematic representation of the *in vivo* intranasal dosing regime, B) Viral lung titres, analysed using TCID_50_, in the supernatant collected from homogenised lung showing a near 4-log (99.99%) reduction in lung viral titres, C) pulmonary damage scores obtained following blind-scoring of hematoxylin and eosin-stained lung tissue sections of antiviral and PBS-treated mice following RSV or sham infection, D-I) Immune cell subset profiling by flow cytometry in RSV-infected and PBS control mice dosed with or without antiviral polymer (see related Suppl. Figure S11 for gating strategy). Data (n = 4-6) is from one representative experiment. Statistics calculated by Kruskal-Wallis test after Shapiro-Wilks normality testing.

Thus, the significant decrease in RSV lung titres clearly shows that **star-PSS** is a highly effective virucidal antiviral *in vivo*. Some caution is required given the polymer causes an increase in some immune cell populations in the lung post-infection, although this did not result in enhanced lung inflammation post-infection, which in fact appeared to be reduced in mice treated with **star-PSS** (Figure 5C). Thus, the combined *in vitro* and *in vivo* efficacy data, together with data showing their low *in vivo* toxicity, negligible effects on the immunological profile and their ease of production, suggest that these star polymers have significant potential to be the first global treatment option for RSV and other viruses.

## Conclusion

Viruses indiscriminately affect humans and animals and are a significant threat to global health and economies. Whilst vaccination is the gold standard for viral prevention, there is a clear need to develop and deploy antiviral interventions that are proactive, rather than reactive, to a viral threat. We have shown that **star-PSS** could be an answer to this global problem.

It is a highly effective broad-spectrum biocompatible virucidal antiviral with potential prophylactic and therapeutic applications. SI values for **star-PSS** are the highest ever reported for an antiviral, highlighting its significant potential. *In silico* studies point to a novel virucidal mechanism, whereby **star-PSS** acts as a molecular glue, between viral surface proteins, exerting a destructive force on the membrane. We show, *via* intranasal administration, that **star-PSS** has minimal toxicity *in vivo*, does not illicit a strong immune response and does not posses strong anti-coagulation properties. This coupled with the *in vivo* results showing highly effective therapy, after just 3 days of treatment, adds to the potential of these materials.

One of the most promising aspects of this technology lies not only in its high virucidal antiviral properties but also in its potential to be an accessible treatment option. **star-PSS** can be produced using readily available and cheap starting materials *via* an environmentally friendly water-based synthetic pathway. The need for low to middle income countries to have access to life changing therapeutics is a matter of public health importance and global development. Further evolution of **star-PSS** and discovery of similar antiviral materials is a promising new avenue of research that will lead to the discovery of novel antivirals with the potential to treat current, and prevent future, viral outbreaks.

## Methods

All solvents and materials used were dry and reaction conditions controlled, with polymerisation carried out under a nitrogen atmosphere and all other reactions carried out in an open atmosphere. Starting materials were sourced from Sigma Aldrich unless otherwise stated.

### Size Exclusion Chromatography (SEC)

All polymers were analysed on an Agilent 1260 Infinity system instrument equipped with a differential refractive index (DRI), viscometry (VS), dual angle light scatter (LS), and variable wavelength UV detectors. The system was equipped with with 2X PL aquagel-OH MIXED-M (400 x 7.5 mm) and a PL aquagel-OH guard column. The eluent was 70% water (pH=9, 0.2 M NaNO_3_) and MeOH 30%. Narrow poly(ethylene oxide) standards were used for calibration between 1,300,0000 - 100 g mol^*-*1^. All samples were filtered through a 0.2 *µ*m nylon filters before injection. The refractive index increment, dn/dc, for PSS in the eluent conditions used was calculated to be 0.154 mL/g.

### ^1^H Nuclear Magnetic Resonance (^1^H NMR)

NMR spectra were recorded on Bruker Avance III HD 400 MHz spectrometer at room temperature using either chloroform-d or deuterated water. The resonance signal of residual solvent at 7.28 and 4.71 ppm respectively served as a reference for the chemical shift, *d*.

### CTA synthesis

Acetone (100 mL), pentaerythritol tetrakis 3-mercaptopropinate (1.0 g, 2.046 mmol), triethylamine (1.65 g, 16.368 mmol), and carbon disulfide (1.620 g, 21.278 mmol) were mixed together and stirred vigorously for 1 hour at room temperature. 2-bromopropanoic acid (1.502 g, 9.821 mmol) was added to the mix and left to stir at room temperature overnight. The reaction was stopped and the mixture taken to pH 1 using hydrochloric acid. The mix was washed against ethyl acetate and DI water and the aqueous layers were combined and collected using a separating funnel before again being reduced and concentrated under reduced pressure. Purification of the material was attained through running an acidified column first with 90% 2:1 hexane to acetone with 10% (v/v) acetic acid, followed by 90% 1:1 hexane to acetone with 10% acetic acid. ^1^H NMR (nuclear magnetic resonance) (400 MHz) spectra were recorded using a Bruker Avance III reference here was CDCl_3_ internal reference set to *d* 7.28ppm. Here the environments of interest are located at ^1^H NMR (CDCl_3_) *d* (ppm): 1.63 (12H, CHCH_3_), 2.81 (8H, COOCH_2_CH_2_), 3.61 (8H, CH_2_CH_2_SC S), 4.14 (8H, COOCH_2_), and 4.80 (4H, CHCH_3_). These matched literature from Dao et al. confirming the desired core NMR.^45^

### Linear Polymer Synthesis

Sodium 4-vinylbenzene sulfonate (1.00 g, 4.85 mmol), linear CTA (4-[[[(2-Carboxyethyl)thio]thioxomethyl]- thio]-4-cyanopentanoic acid) (29.82 mg 0.097 mmol), and 4,4’-Azobis(4-cyanopentanoic acid) (2.72 mg 0.0097 mmol) were dissolved in 4mL of deionised water and nitrogen degassed for 20 minutes while stirring. The reaction was held under nitrogen and kept at 70^*o*^C until polymerisation was at approximately 96% conversion (4 hours). The reaction was temperature and oxygen quenched and dialysed using a 1kDa pore membrane in deionised water for 3 days and two changes per day.

### General Star Polymer Synthesis

The following methodology outlines the procedure for synthesising one gram of Polystyrene Sulfonate (PSS) Polymer four arm star with a degree of polymerisation (DP) 50. sodium 4-vinylbenzene sulfonate (1 g, 4.85 mmol), 4 arm CTA (26.22 mg 0.024 mmol), and 4,4’-Azobis(4-cyanopentanoic acid) (0.67 mg 0.0024 mmol) were dissolved in 4mL of 0.1M sodium hydroxide in deionised water and nitrogen degassed for 20 minutes while stirring. The reaction was held under nitrogen and kept at 65^*o*^C until polymerisation was at approximately 90% conversion (16 hours). The reaction was temperature and oxygen quenched and dialysed in deionised water for three days and two changes per day. Polymers were dried under reduced pressure to yield a pale yellow film or freeze-dried to yield a very pale yellow fluffy powder. ^1^H NMR (400 MHz) spectra were recorded using a Bruker Avance III reference here was D_2_O internal reference set to *d* = 4.71ppm. Broad peaks were found at 1.36ppm, 6.56ppm, and 7.45ppm (Figure S2), yield was calculated pre-dialysis as 90% conversion with each 1g batch yielding around 800-850mg of polymer.

### Molecular Dynamic Simulations

Nanoscale Molecular Dynamics^75^ simulations were performed on the crystallographic structures of RSV (PDB: 5UDE). All the systems are modelled with CHARMM36 protein force field^76^. The cutoffs of van der Waals and Coulombic interactions were 10 Å, and the long-range Coulombic interactions were calculated by the PME method (periodic boundary conditions)^77^. The simulations were performed in the NPT ensemble (p = 1 bar and T = 310 K), using the Langevin dynamics with a damping constant of 1 ps-1, and a time step of 2 fs. After 2 ns of minimization, the systems were simulated up to 200 ns, where we applied a full protein constraints for the single protein-polymer simulations.

For the assemblies simulations, membrane constraints were applied where 3 atoms at the bottom for each sub-units where held in harmonic potential (exponent for the harmonic constraint energy function 2 and force constant k 0.5). The systems were simulated for 350 ns at 298K. For this simulation the non-bonded energies were calculated by considering the 3 closest proteins to one polymer. The distances were calculated by averaging the center of the mass over the time of 3 proteins close to the previous polymer. The force was then calculated by calculating the slope of the best fit line of energies over the distance. The hydrogen bonds were evaluated with the HBonds Plugin in VMD and the default setup (cutoff distance of 3 Å, cutoff angle of 20°)^78^.

### Material Qualification

Cleavage of the star to the proton began by weighing out reagents in quantities proportional to the theoretical arm lengths of each polymer in question. Per 0.5g of polymer in question was added to a vial followed by: 17.9mg of 1-Ethylpiperidine hypophosphite, 0.66mg of 2,2’-Azobis(2-methylpropionitrile), along with 2.5mL of N,N-Dimethylformamide, the mixture then degassed under nitrogen for 20 mins The vial was then heated in an oil bath with continuous stirring at 100^*o*^C for 3 hours. Following this the mix will be precipitated in hexane and the sludge freeze-dried before being re-solubalised in GPC reagents and run through the machine. The resulting molecular weights of the cleaved material versus the normal shows if the configuration of the polymer is truly star shaped as well as if the arm lengths of the polymer lines up with the expected.

### Biological Testing

All biological testing was conducted in accordance with standardised testing procedures. Reagents sourced from ThermoFisher Scientific unless otherwise stated and used as purchased unless specific modification mentioned. All reagents and materials were sterile and handled using sterile techniques, work was carried out in class II biological safety cabinets. Each test was conducted in at least duplicate and triplicate where possible. Analysis was performed using GraphPad Prism (California) Version 9.1.0 (216).

#### Cells

Cells sourced from ATCC were; Vero CCL-81, and HeLa derived HEp-2 cells. Additionally Mv1-Lu cells were gifted by Professor Pamela Vallely from the Faculty of Biology, Medicine and Health (FBMH) at the University of Manchester. Cells were propagated in appropriate growth media; Dulbecco’s modified Eagle’s medium (DMEM) supplemented with 10% heat-inactivated Foetal Bovine Serum (FBS) and 1% penicillin/streptomycin (Sigma Aldrich). Cells were incubated throughout at 37^*o*^C with 5% CO_2_.

Where cell and viral culture relating to the FAIRY assay is concerned, all cells and viruses were grown in Dulbecco’s Modified Eagle Medium (DMEM) modified with high glucose, L-glutamine, sodium pyruvate but the absence of phenol red. Media was supplemented with 1% penicillin/streptomycin only *i*.*e*. no addition of heat-inactivated Foetal Bovine Serum (FBS).

#### Viruses

Viruses were either purchased from ATCC or gifted by Professor Pamela Vallely from the Faculty of Biology, Medicine and Health (FBMH) at the University of Manchester. Viruses donated were HSV-2, RSV, HCoV OC43, and CMV. HRV8 was sourced from ATCC. Viral stocks were kept at -80^*o*^C and each thaw recorded due to inevitable decrease in virus concentration with thaw cycles.

#### Viral Inhibition Assays

Three main inhibition assays were dose response assays, virucidal assays, and median tissue culture infectious dose assays (TCID_50_). Reagents for these assays included phosphate-buffered Saline (PBS), DMEM with 2% FBS, Methylcelloulose (MTC), and Crystal Violet. Methylcellulose was prepared by adding 3g of MTC to 200mL of deionised water before autoclaving overnight, 60mL of this stock was then added to 140mL of FBS-free DMEM before being stirred at room temperature for 15 mins then stored at 4^*o*^C. Before use MTC must be stirred at room temperature for 2 hours. Crystal violet solutions were prepared for use at 0.1% crystal violet in 79.9% water and 20% ethanol.

#### Dose Response Assay

These assays began with seeding cells into a 24 well plate and incubating until 90% confluency (virus dependent as certain viruses grow slower and therefore the initial cell numbers would be seeded at a lower concentration). Stock concentrations of the antiviral material were diluted down from 500*µ*g/mL to 0.5ng/mL in PBS prior to experiment and stored at 4^*o*^C for use throughout. Virus stocks were made by diluting 30*µ*L of virus into 570*µ*L for each antiviral concentration and allowed to incubate together for 1 hour at 37^*o*^C. The media on the cells was replaced with the antiviral material and virus solution and allowed to incubate for a further hour. The antiviral virus mix was finally replaced with MTC and incubated to allow for visible plaques to form. Following this, MTC was replaced with enough crystal violet stain (described previously) to cover and left to stain for 30 mins. Samples were washed 3x in deionised water and left to dry overnight, finally plaques were counted using a light microscope. Internal controls included a non-treatment control showing viral infection without any intervention.

#### Virucidal Assay

Virucidal assays began with seeding cells into a 96 well plate until 90% confluency (again virus dependent). Viral stocks were incubated in the antiviral material half and half for an hour at 37^*o*^C, non treatment controls were viral stock and PBS only. Antiviral and virus stock was titrated 1:10, 3 times before being added to the confluent cells and further titrated down the plate in triplicate. Finally in the same manner as before the supernatant was removed and replaced with MTC, and allowed to incubate at 37^*o*^C until visible plaques formed and then subsequently stained with crystal violet, dried, and counted as described previously. Internal controls included a non-treatment control showing viral infection without any intervention.

#### TCID_50_ Assay

TCID_50_ assays began with seeding cells into 96 well plates (10,000 cells per well) and allowing to attach for at least 4 hours. Samples were added to the plates in quadruplicate. Each of the first four wells in the first column of the plate had 180 *µ*L of media added to them, the rest of the plate had 100 *µ*L of media added. 20 *µ*L of sample was added into the first four wells in column 1 (A1-D1) and mixed by pipetting. 100 *µ*L was taken from the first column and titrated across towards the end of the plate (wells A11-D11). The 100 *µ*L titration was continued from wells A12-D12 to E1-H1 and across to E12-H12. The last four wells in column 12 (A12-D12) were left as negative controls without virus added. The plates were then left until cytopatic effect was observed using a light microscope with a 10x objective. Viral titer was calculated using the Spearman and Kabërmethod.

#### Fluorescence Assay for vIRal IntegritY (FAIRY)

FAIRY serves as a nucleic acid fluorescent readout for the infectivity of a virus following treatment, as previously described.^52^ Briefly, PSS antiviral and first thaw HSV-2, RSV, OC43 and HRV-8 stocks in phenol red-free DMEM (with no FBS supplemented) were incubated at a 1:1 ratio for 1 hr at 37*o*C. The non-treated sample (NTC) was incubated with PBS, whilst the blank controls were medium only. Subsequently, 50 *µ*L of TO-PEG was added to 50 *µ*L of sample into a black 96-well flat bottom plate. Following a 10 min incubation, fluorescent readouts were measured using Envision ^®^ multimode monochromator plate reader by PerkinElmer Inc., (V1.13.3009.1394) (*l* _*Ex*_=510 nm and *l* _*Em*_=533 nm)

#### Cytotoxicity Assay

Cytotoxicity testing was conducted in order to assess the biocompatibility of the antiviral material being researched. Using the standard protocols provided by the manufacturer the assay was conducted in order to assess the cytotoxicity of the synthesised material. Briefly the methodology was as follows. Cells were seeded into 96 well plates to 90% confluency, a further plate was prepared containing media only as a background control. High concentrations of 50 *µ*L of polymer in PBS were added to each well in triplicate for a final volume of 100 *µ*L, starting with 5 mg/mL and continuing with 2.5 and 1 mg/mL and followed by 500, 250, 100, and finally 50 *µ*g/mL, control plate had PBS without polymer added into it, and control wells with cells and PBS were prepared as the control. Plates were allowed to incubate for 24 hours at 3^*o*^C with 5% CO_2_. After incubation 20 *µ*L of MTS reagent (CellTiter 96® Aqueous One Solution Reagent) was added to each well and allowed to incubate for 4 hours at 37^*o*^C. Finally the plates were read at 490nm absorbance and recorded for cytotoxicity.

### Thromboelastography

Investigation of the effect of linear PSS and **star-PSS** polymers on clot forming ability was conducted using a TEG5000 thromboelastograph (Haemonetics SA, Signy, Switzerland). Controls for the clotting behaviour of normal blood (Level I Biological Control) and hypercoagulable blood, or blood that tends towards increased clotting (Level II Biological Control) were also purchased from Haemonetics SA and prepared as indicated on the packaging. Each ampule of lyophilised control material contained 2 volumes of clotting material, which were always run in parallel (TEG5000 is a 2-pin instrument) and then repeated three times from different batches of material. Lyophilised controls were allowed to warm to room temperature for 10 min, and then 1 ml ultrapure water was added. Samples were shaken to assist dilution, allowed to sit for 5 min, then shaken again and allowed to rest for a further 5 min. Controls containing no polymer were prepared for measurement by pipetting 20 *µ*L of a 20 mM solution of CaCl_2_ into the measurement cup, followed by 340 *µ*L of either the Level I or Level II control. The measurement pin was immediately inserted into the cup and the measurement started.

To measure the effect of **star-PSS** polymers, linear PSS or heparin sulfate on the coagulation ability of the clotting controls, solutions of 0.06 mg/ml and 0.1 mg/mL of each of the polymers were prepared in 20 mM CaCl_2_. This was done to ensure no dilution of the clotting materials by the further addition of polymer solutions to the mixture. 20 *µ*L of a solution containing the polymer of interest in 20 mM CaCl_2_ was then pipetted into the measurement cup, followed by 340 *µ*L of either the Level I or Level II control and measured immediately. All polymers were tested 3 times, with the exception of heparin, as at both concentrations no clotting was detected by the instrument, it was decided a third replicate was unnecessary.

### in vivo intranasal toxicity and associated histology

Mice (C57BL/6, Charles river) were housed at an AALAC International-accredited institution and ethically cared for in accordance with The Guide for the Care and Use of Laboratory Animals. Mice (n = 5, C5BL/6, female) were dosed with 100 *µ*g **star-PSS** each day for 5 days, administered intranasally (10 *µ*L of 1 wt.% antiviral polymer in phosphate-buffered saline). Control group mice (n = 4) received a vehicle control, administered intranasally (10 *µ*L PBS) each day for 5 days. Mouse weights were recorded each day. At the end of the 5 days, blood was collected from all specimens for serum biochemistry toxicology measurements and histology was performed on heart, liver, kidney, and nasal turbinates. Heart, liver, and kidneys were immersion-fixed in 10% neutral buffered for 72 hours. The skull and nasal cavity were fixed in Cal-Ex II Fixative/Decalcifer (Fisher Scientific) for 72 hours. Following decalcification, 4 transverse sections (2mm thick) of the nasal cavity were obtained for evaluation.

### Intranasal RSV infection and antiviral polymer dosing

Mice were anaesthetized by inhalation isoflurane and infected intranasally with 1.6×10^6^ PFU of RSV or PBS sham in a volume of 30*µ*L PBS on day 0. On days 1 - 3 post-infection, mice were intranasally dosed with 25*µ*g antiviral polymer (PSS DP50 4-Arm) or PBS sham in a volume of 20*µ*L. Daily weights were monitored until day 4 post-RSV infection.

### Tissue harvest and processing for flow cytometry, viral load and histology

Mice were culled on day 4 post-RSV infection, lungs and spleen were collected after PBS perfusion with 10mL of PBS-1mM EDTA. For flow cytometry, one lobe of lung and half of spleen tissue, were first weighed. Tissues were then minced and digested with either collagenase type I (10mg/mL) and DNAse I (50*µ*g/mL) for 40 minutes (lungs), or with collagenase type D (1mg/*µ*L) and DNAse I (50*µ*g/mL) for 30 minutes (spleens), at 37 degrees in a shaking incubator. The tissue digest was passed through a 70 micron filter, washed with 25mL of cold PBS-5mM EDTA solution, spun down at 500g and RBCs lysed with a commercial RBC lysis buffer. After 1x PBS wash, the single cell suspension was resuspended in complete RPMI media for downstream procedures. For viral load assay, three lung lobes were collected into sterile PBS in RINO homogenizer tubes with a pre-loaded stainless steel bead. Lung tissues were weighed, and homogenized by bead-milling (2x times for 2 minutes on medium speed), or until homogenate does not contain tissue chunks. Viral titres were then determined using a TCID_50_ assay, specifically, lung lysate was added to pre-plated cells (10,000 cells per well) that were given 4 hours to attach. Samples were added to the plates in quadruplicate. The first four wells in the first column of each plate had 180 *µ*L of media added to them, the rest of the plate had 100 *µ*L of media added. 20 *µ*L of sample was added into the first four wells in column 1 (A1-D1) and mixed by pipetting. 100 *µ*L was taken from the first column and titrated across towards the end of the plate (wells A11-D11). The 100 *µ*L titration was continued from wells A12-D12 to E1-H1 and across to E12-H12. The last four wells in column 12 (A12-D12) were left as negative controls without virus added. The plates were then left until CPE was observed using a light microscope with a 10x objective. Viral titre was calculated using the Spearman and Kabër method. For histology, the left lung lobe was inflated with 2mL of 10% formalin, the spleen was halved longitudinally and both tissues were fixed in 10% formalin for 24h. Formalin-fixed tissues were then transferred to 70% EtOH solution before paraffin wax embedding. Samples were prepared for histology analysis using a standard hematoxylin and eosin staining protocol. Samples were embedded in paraffin wax and sectioned several times to reveal a consistent surface. Sections were loaded onto slides and dewaxed using xylene at 65 ^*o*^C, samples were dehydrated in three washes of degrading ethanol and washed with water. Slides were then bathed in Haematoxylin for 2 minutes followed by a washing regime in 5% Acetic acid followed by water, Scotts Tap Water / Bluing for 30 seconds and water again. The final staining was conducted in Eosin for 1 minute and 30 seconds and finally washed again in ethanol followed by water, ethanol again, and finally 2 washes of xylene for 2 minutes each.

### Flow cytometry and antibody staining

Single cell suspensions (all of lung and 1/5th of spleen) was washed 1x with PBS, spun down and resuspended in 300*µ*L of Zombie UV Live/Dead stain for 15 minutes. Cells were washed with 1x PBS and resuspended in 50*µ*L of Fc blocking solution in FACS buffer (2% FCS in PBS-2mM EDTA). After 10 minutes, 10*µ*L of antibody staining mastermix (see table S2) prepared in BD Brilliant Stain Buffer Plus, was added to lung and spleen samples in Fc Block and incubated for 30 minutes at 4 degrees, protected from light. The antibody mastermix was washed off with 1x PBS, and cells were fixed in 300*µ*L of fixation/permeabilization buffer for 20 minutes. The fixed samples were washed with 3mL of 1x PBS, passed through a 35 micron filter, and resuspended in 300*µ*L of PBS. 25*µ*L of counting beads were spiked into each sample before flow cytometry acquisition, to calculate for cell numbers per 1g of tissue in downstream data analysis.

### Histopathology

Histopathology images were gathered through the imaging service at the University of Manchester. Images were gathered at 40x brightfield using the 3D Histech Pannoramic P250. Images were then analysed using the 3D Histech slideviewer software by a board-certified veterinary pathologist (KMC). Pulmonary damage was assessed using an ordinal (0-4) grading scheme for the following criteria: alveolar septal necrosis, bronchiolar degeneration and regeneration, early interstitial fibrosis, interstitial and alveolar pneumonia (mixed infiltrates), and perivascular/peribronchiolar lymphoplasmacytic inflammation. Each criterion was graded based on the percentage of the pulmonary parenchyma affected (0 = absent, 1 = minimal (<5% of parenchyma), 2 = mild (5-25% of parenchyma), 3 = moderate (25-50% of parenchyma), 4 = severe (>50% of parenchyma)). A cumulate pulmonary damage score was calculated for each mouse as a sum of the individual criteria.

## Supporting information

ESI

## Acknowledgements (not compulsory)

Diagnostic Laboratory, Stanford University, Dept. of Comparative Medicine performed serum biochemistries. L.J.B acknowledges the Biotechnology and Biological Sciences Research Council and the Manchester Doctoral Training Programme through Grant DTP3 2020-2025, Reference BB/T008725/1

## Author contributions statement

E.H.S. synthesised and characterised star-PSS and L-PSS, performed *in vitro* assays on all viruses, prepared histology samples for imaging and supported *in vivo* assays. S.M.L. conducted *in vivo* RSV model and prepared and ran all FACS analysis. U.C.C. supported *in vivo* RSV model and associated data collection. F.C. performed all MD simulations. O.S. coordinated *in vivo* toxicity study and administered intranasal doses. Y.E.S. supported *in vivo* toxicity study. K.M.C. performed all histopathology analysis. L.J.B performed the FAIRY assay against RSV, OC43, HRV8 and CMV. S-L.M. developed the FAIRY assay and demonstrated its capabilities with the STAR polymer using HSV-2. R.H.T.B. supported polymer synthesis and characterisation. Z.W.K. performed all blood coagulation studies and associated analysis. P.K. coordinated all MD simulations. E.A.A. oversaw *in vivo* toxicity studies and associated data analysis. M.A.T. coordinated and supported *in vivo* RSV studies and associated analysis. S.T.J. conceptualised polymer antivirals, conducted initial synthesis and testing, oversaw all aspects of the project and its management and wrote the manuscript. All authors reviewed the manuscript.

## Additional information

The authors declare no competing interests.

## Notes

### Competing Interest Statement

The authors have declared no competing interest.

