## Supplementary material for "Star-polymers as potent broad-spectrum extracellular virucidal antivirals": ESI

[1]Department of Materials, The University of Manchester, Manchester, M13 9PL, UK
[2]The Henry Royce Institute, Manchester, M13 9PL, UK
[3]Faculty of Biology, Medicine and Health, University of Manchester, Manchester, M13 9PL, UK
[4]Department of Materials Science & Engineering, Stanford University, CA, 94305, USA
[5]Department of Comparative Medicine, Stanford University School of Medicine, CA, 94305, USA
[6]Department of Chemistry, University of Basel, 4058 Basel, Switzerland
[7]Department of Chemistry, University of Illinois at Chicago, Chicago, Illinois 60607, United States
[8]Department of Physics and Department of Biopharmaceutical Sciences, University of Illinois at Chicago, Chicago, Illinois 60607, United States
[9]School of Chemistry, University of Birmingham, Edgbaston, Birmingham, B15 2TT
[*]

### ABSTRACT

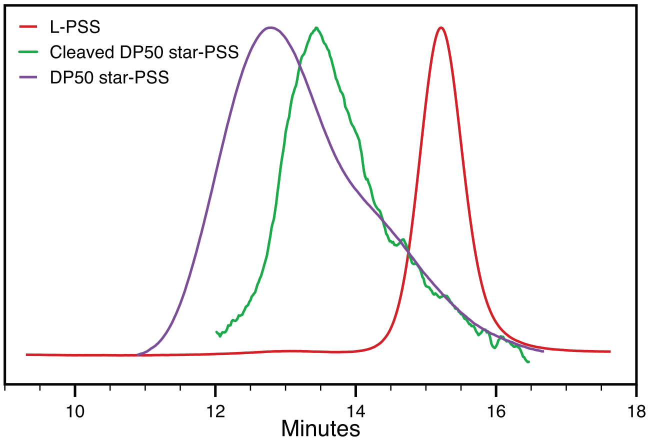


| Sample | $M_n$ | MW | Dispersity |
|---|---|---|---|
| L-PSS | 2776 | 4358 | 1.57 |
| Cleaved star-PSS | 20901 | 36286 | 1.74 |
| star-PSS | 27297 | 50196 | 1.84 |

**Fig. S1.** Gel permeation chromatogram for **L-PSS**, **star-PSS** and cleaved **star-PSS** and associated table containing $M_n$, MW and dispersities for each polymer.

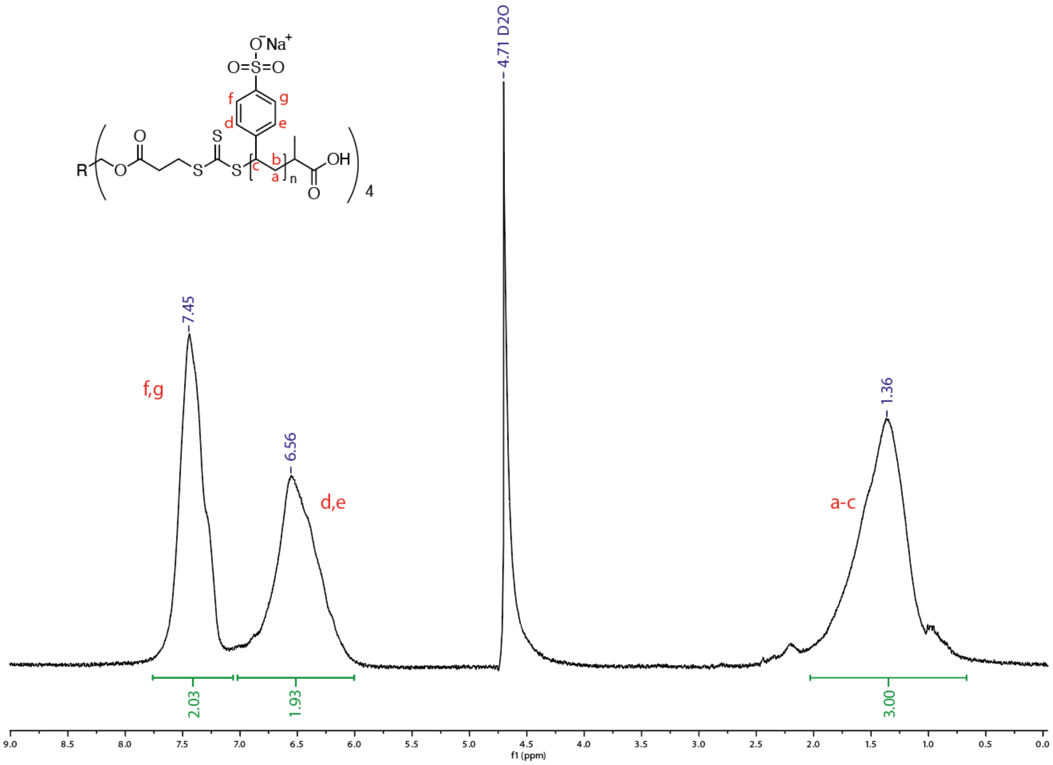


**Fig. S2.** NMR spectra of **star-PSS** in $D_2O$.

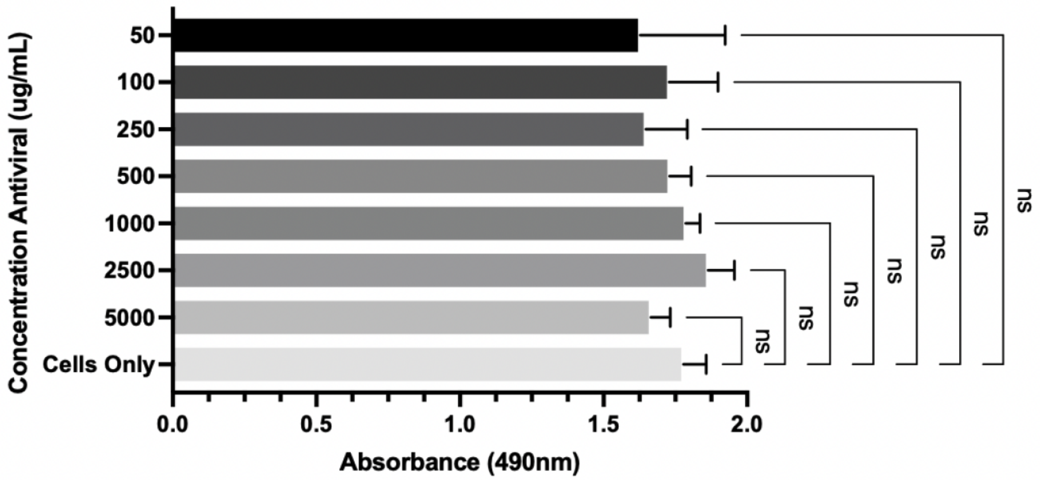


**Fig. S3.** MTT assay up to 5mg of **star-PSS** in direct contact with cells showing no significant difference in absorbance.

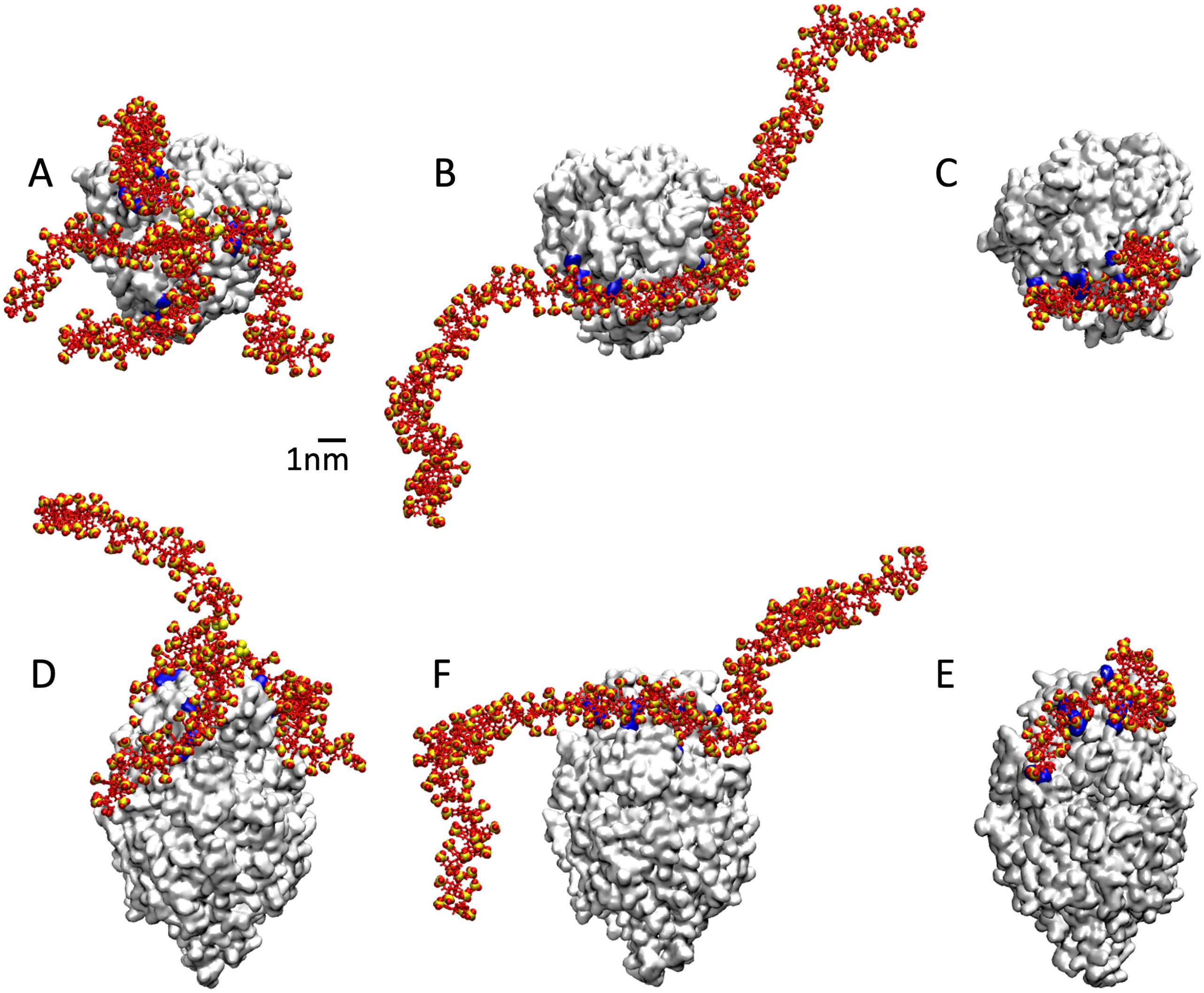


**Fig. S4.** Single protein simulations for RSV with the 3 polymers. (A-C) top view, for PSS star-like, linear 200 and linear 50, (D-E) the side view. On the protein surface (white) we show the basic residues in contact with the polymer.

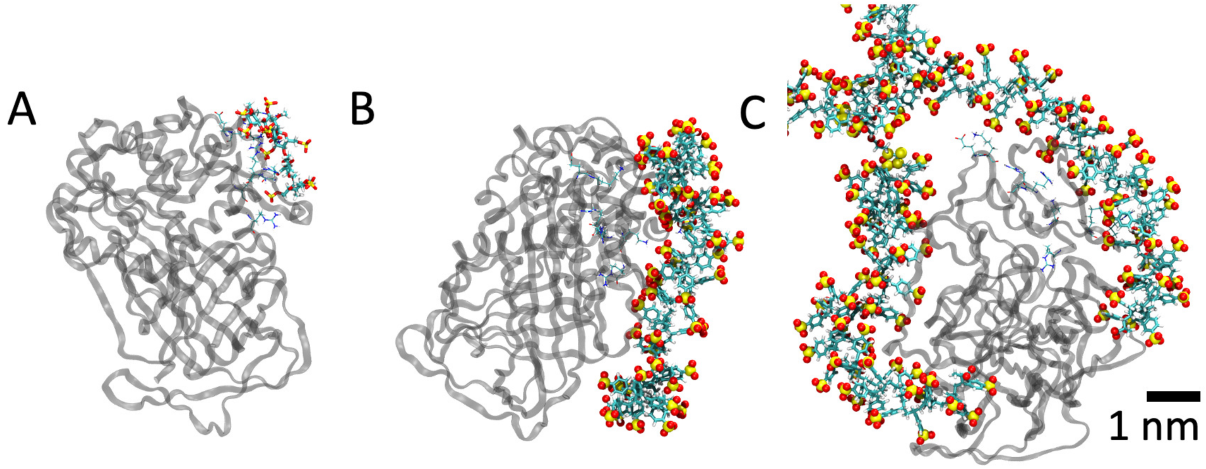


**Fig. S5.** Antithrombin interacting with (A) heparin, (B) linear PSS and (C) star-like PSS.

| resid | resname | Heparin (MAX 106) | Star-like (MAX 182) | Linear (MAX 225) |
|---|---|---|---|---|
| 11 | LYS | 0.84 | 1.00 | 1.00 |
| 13 | ARG | 0.99 | 0.91 | 0.97 |
| 46 | ARG | 1.01 | 0.00 | 0.47 |
| 47 | ARG | 1.01 | 0.00 | 0.47 |
| 113 | GLU | 1.01 | 0.36 | 0.32 |
| 114 | LYS | 1.01 | 0.93 | 0.62 |
| 125 | LYS | 0.99 | 0.87 | 0.46 |
| 129 | ARG | 1.00 | 0.00 | 0.08 |
| **Energy (kcal/mol)** | | **-70.76** | **-181.43** | **-93.13** |

**Fig. S6.** Contact over the time over with specific residue important for the heparin-antithrombin interaction. The values were normalized over the trajectory length. We also reported in the last raw the non-bonded energies.

| Heparin | | Star-like | | Linear 200 | |
|---|---|---|---|---|---|
| SegPROB-LYS28-Side | 0.50% | SegPROB-LYS11-Side | 27.50% | SegPROB-LYS11-Side | 23.50% |
| SegPROB-ARG46-Side | 34.50% | SegPROB-ALA30-Main | 0.50% | SegPROB-ARG129-Side | 0.50% |
| SegPROB-ARG47-Side | 39.00% | SegPROB-LYS29-Side | 36.00% | SegPROB-ARG13-Side | 48.50% |
| SegPROB-ASN45-Main | 40.00% | SegPROB-ARG24-Side | 64.00% | SegPROB-LYS114-Side | 15.00% |
| SegPROB-ARG47-Side | 65.50% | SegPROB-ASN45-Side | 0.50% | SegPROB-LYS28-Side | 20.00% |
| SegPROB-LYS114-Side | 51.50% | SegPROB-LYS114-Side | 15.50% | SegPROB-LYS136-Main | 15.50% |
| SegPROB-ASN45-Side | 28.50% | SegPROB-ALA293-Main | 1.00% | SegPROB-ASN135-Main | 4.50% |
| SegPROB-ASN45-Side | 22.00% | SegPROB-LYS28-Side | 19.50% | SegPROB-THR31-Side | 7.50% |
| SegPROB-LYS11-Side | 17.50% | SegPROB-LYS294-Side | 14.50% | SegPROB-SER137-Side | 14.50% |
| SegPROB-ARG13-Side | 28.00% | SegPROB-ARG235-Side | 2.50% | SegPROB-ARG13-Main | 8.50% |
| SegPROB-ARG13-Main | 4.50% | SegPROB-ARG13-Side | 36.50% | SegPROB-LYS275-Side | 23.00% |
| SegPROB-GLU113-Main | 25.50% | SegPROB-LYS290-Side | 2.00% | SegPROB-LYS136-Side | 31.50% |
| SegPROB-LYS114-Main | 42.00% | SegPROB-THR9-Side | 37.00% | SegPROB-ARG132-Side | 16.50% |
| SegPROB-LYS114-Side | 8.50% | SegPROB-LYS125-Side | 14.50% | SegPROB-VAL5-Main | 3.50% |
| SegPROB-ARG46-Side | 19.50% | SegPROB-LEU292-Main | 0.50% | SegPROB-LYS29-Main | 26.50% |
| SegPROB-LYS125-Side | 38.50% | SegPROB-LYS290-Side | 20.50% | SegPROB-THR31-Main | 6.00% |
| SegPROB-LYS125-Side | 24.50% | SegPROB-ARG406-Side | 10.00% | SegPROB-SER230-Side | 0.50% |
| SegPROB-GLN118-Side | 0.50% | SegPROB-LYS287-Side | 37.50% | SegPROB-SER227-Side | 2.00% |
| SegPROB-ARG129-Side | 17.50% | SegPROB-TYR240-Side | 0.50% | SegPROB-SER25-Main | 0.50% |
| SegPROB-ARG13-Side | 26.50% | SegPROB-TYR260-Side | 12.50% | SegPROB-ASN135-Side | 2.00% |
| SegPROB-ARG13-Side | 1.00% | SegPROB-ARG262-Side | 51.00% | SegPROB-LYS226-Side | 8.00% |
| SegPROB-LYS11-Side | 9.00% | SegPROB-LYS107-Side | 17.00% | SegPROB-LYS228-Side | 15.50% |
| SegPROB-THR44-Side | 3.50% | SegPROB-LYS114-Main | 0.50% | SegPROB-LYS222-Side | 1.00% |
| SegPROB-SER112-Side | 1.50% | SegPROB-ASN405-Side | 3.50% | SegPROB-LYS139-Side | 17.50% |
| SegPROB-ALA43-Main | 0.50% | SegPROB-PHE106-Main | 1.50% | SegPROB-ARG47-Side | 17.50% |
| SegPROB-ARG46-Main | 0.50% | SegPROB-SER191-Side | 20.00% | SegPROB-ARG46-Side | 7.50% |
| | | SegPROB-LYS136-Side | 23.00% | SegPROB-LYS114-Main | 1.00% |
| | | SegPROB-VAL190-Main | 13.00% | SegPROB-ASP33-Main | 1.50% |
| | | SegPROB-SER191-Main | 0.50% | SegPROB-LYS28-Main | 7.00% |
| | | SegPROB-LYS294-Side | 33.50% | SegPROB-GLU27-Main | 4.00% |
| | | SegPROB-LYS169-Side | 2.50% | SegPROB-LYS133-Main | 1.00% |
| | | SegPROB-LYS139-Side | 11.00% | SegPROB-LYS125-Side | 2.50% |
| | | SegPROB-LYS193-Side | 2.50% | SegPROB-ALA30-Main | 0.50% |
| | | SegPROB-LYS348-Side | 2.00% | SegPROB-TYR220-Side | 0.50% |
| | | SegPROB-ARG57-Side | 18.50% | SegPROB-GLU32-Main | 1.00% |
| | | SegPROB-LYS297-Side | 8.00% | SegPROB-SER138-Main | 2.00% |
| | | SegPROB-GLU296-Main | 6.50% | SegPROB-LYS193-Side | 3.50% |
| | | SegPROB-ARG132-Side | 0.50% | SegPROB-ASP6-Main | 5.00% |
| | | SegPROB-ASP14-Main | 1.00% | SegPROB-THR9-Side | 8.00% |
| | | SegPROB-LYS29-Main | 1.00% | SegPROB-LYS29-Side | 0.50% |
| | | SegPROB-LYS241-Side | 1.50% | SegPROB-SER138-Side | 0.50% |
| | | SegPROB-GLU289-Main | 1.00% | SegPROB-LYS11-Main | 2.50% |
| | | SegPROB-THR9-Main | 1.00% | SegPROB-SER137-Main | 1.00% |
| | | SegPROB-HSD65-Side | 2.00% | SegPROB-GLU113-Main | 0.50% |
| | | SegPROB-SER56-Side | 7.50% | SegPROB-LYS257-Main | 4.50% |
| | | SegPROB-ASN127-Side | 0.50% | SegPROB-LYS257-Side | 5.50% |
| | | SegPROB-TRP189-Side | 2.00% | SegPROB-LYS228-Main | 0.50% |
| | | SegPROB-LYS53-Side | 2.00% | | |
| | | SegPROB-ALA242-Main | 1.00% | | |

**Fig. S7.** Hydrogen bonds developed from Heparin and polymers with the antitrombin.

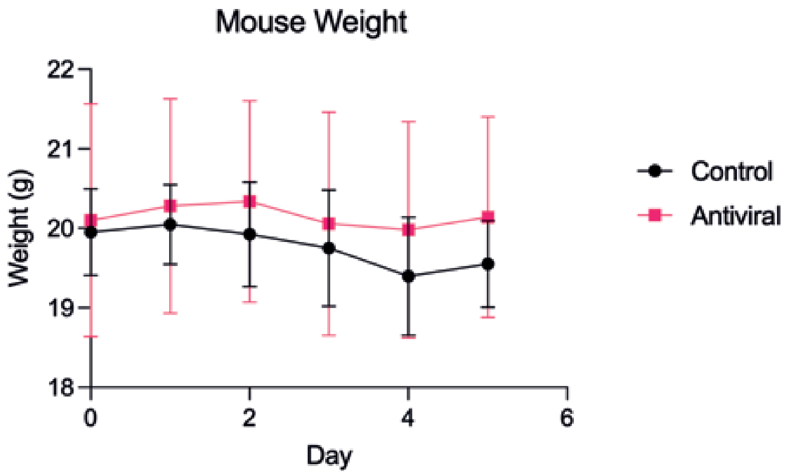


**Fig. S8.** Mouse weights during intranasal dosing of 100$\mu$g once a day for 5 days.

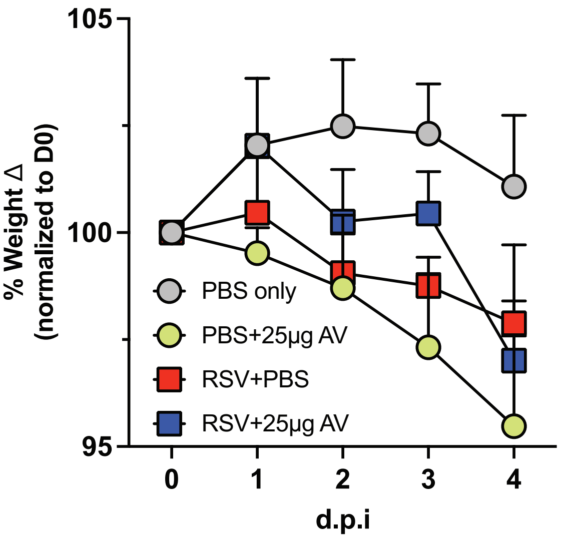


**Fig. S9.** Weight loss curve of PBS only (grey circles), PBS + 25$\mu$g antiviral (green circles), RSV-infected (red squares) and RSV-infected + 25$\mu$g antiviral (blue squares) groups, relative to D0 weights.

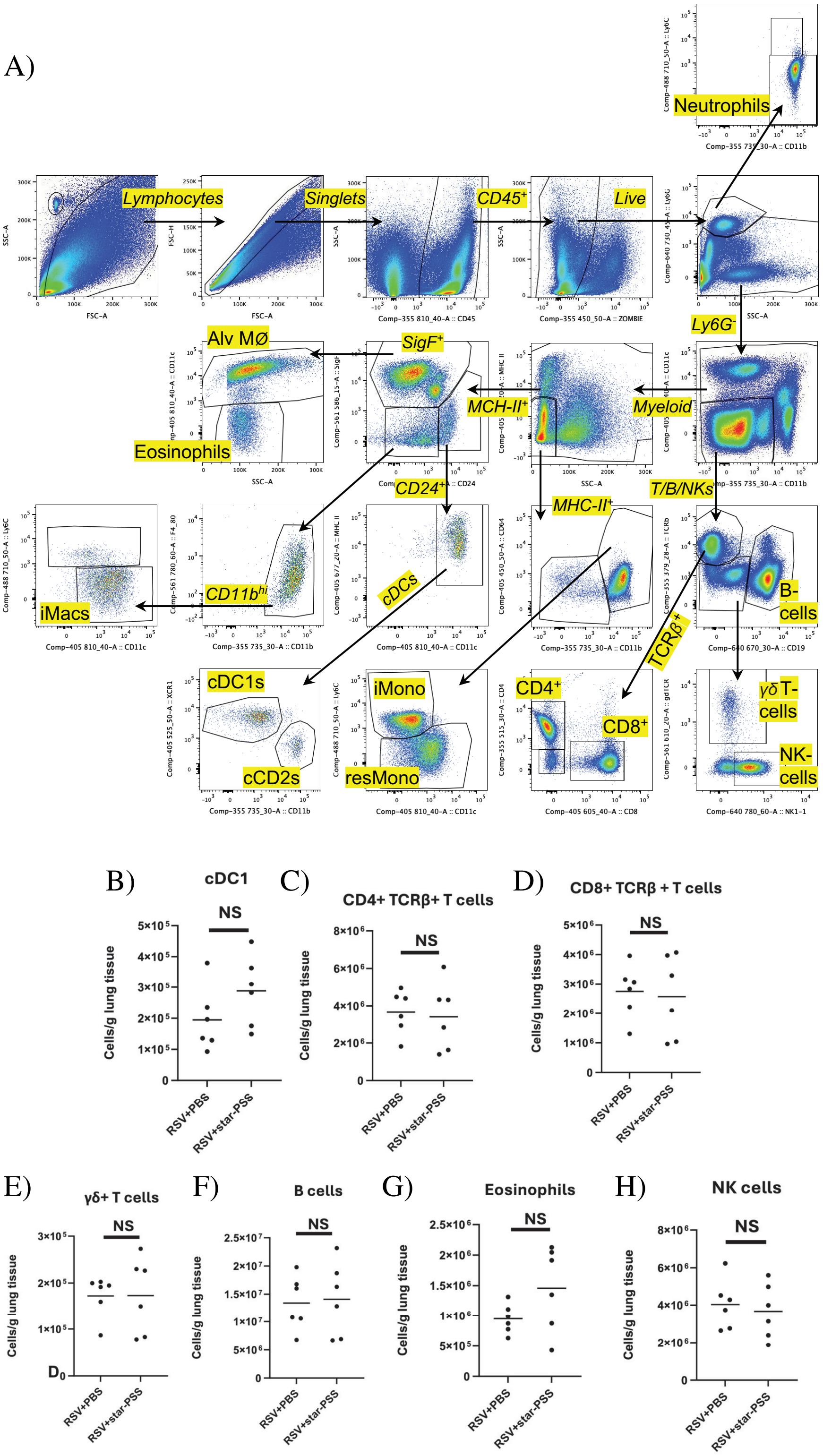


**Fig. S10.** A) Representative flow plots showing the flow cytometry gating strategy used to profile individual immune cell subsets during RSV or sham infection, B-K) Collated data with the cell numbers per 1g of lung tissue for major immune cell subsets profiled in antiviral or PBS-treated mice, 4 days following during RSV or sham infection. Data (n=4-6) is from one representative experiment. All data sets tested for normality by Shapiro-Wilks, if both data sets passed normality test an unpaired t-test done, if not, a Mann-Whitney test done.

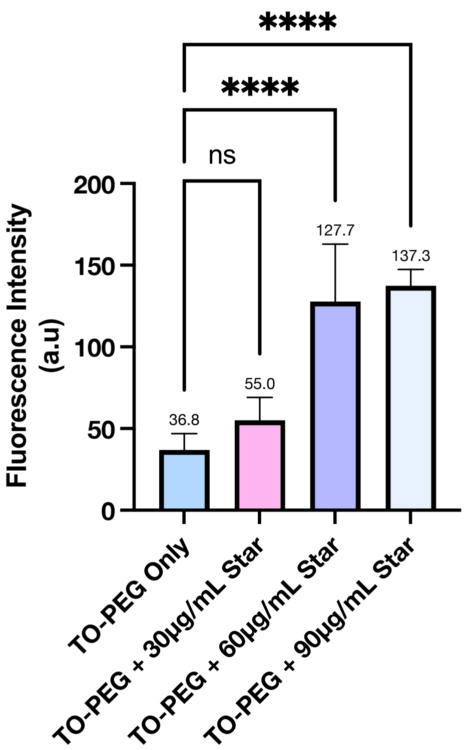


**Fig. S11.** Controls for the FAIRY assay conducted in Figure 2D, showing the effect of increasing polymer concentration on the fluorescence readout of the assay.

| Reagent | Company | Catalog No. |
|---|---|---|
| Ethylenedinitrilo)tetraacetic acid disodium salt | Sigma Aldrich, Merck | 03690-100ML |
| Collagenase, Type I | Gibco,ThermoFisher Scientific | 17100017 |
| Collagenase D | Roche, Merck | 11088858001 |
| Deoxyribonuclease I | Sigma Aldrich, Merck | D4513-1VL |
| DNase I | Roche, Merck | 11284932001 |
| Red Blood Cell Lysing Buffer Hybri-Max | Sigma Aldrich, Merck | R7757-100ML |
| Formalin solution, neutral buffered, 10% | Sigma Aldrich, Merck | HT501128-4L |
| BD Horizon™ Brilliant Stain Buffer Plus | BD Biosciences | 566385 |
| CountBright™ Absolute Counting Beads | LifeTechnologies, ThermoFisher Scientific | C36950 |
| eBioscience™ Foxp3 / Transcription Factor Fixation/Permeabilization Concentrate and Diluent | Invitrogen, ThermoFisher Scientific | 00-5521-00 |

**Table S11.** List of experimental reagents

| Antibody Marker / Conjugate | Clone | Company | Catalog No. |
|---|---|---|---|
| Zombie UV™ Fixable Viability Kit | | BioLegend | 423108 |
| BD Pharmingen™ Purified Rat Anti-Mouse CD16/CD32 | 2.4G2 | BD Biosciences | 553142 |
| APC anti-mouse CD19 | 6D5 | BioLegend | 115512 |
| Alexa Fluor® 700 anti-mouse Ly-6G | 1A8 | BioLegend | 127622 |
| Brilliant Violet 711™ anti-mouse CD24 | M1/69 | BioLegend | 101851 |
| Brilliant Violet 785™ anti-mouse CD11c | N418 | BioLegend | 117336 |
| Brilliant Violet 650™ anti-mouse I-A/I-E | M5/114.15.2 | BioLegend | 107641 |
| Brilliant Violet 421™ anti-mouse CD64 (FcRI) | W18349C | BioLegend | 164407 |
| Brilliant Violet 510™ anti-mouse/rat XCR1 | ZET | BioLegend | 148218 |
| APC/Cyanine7 anti-mouse NK-1.1 | PK136 | BioLegend | 108724 |
| PE/Cyanine7 anti-mouse F4/80 | BM8 | BioLegend | 123114 |
| PerCP/Cyanine5.5 anti-mouse Ly-6C | HK1.4 | BioLegend | 128012 |
| PE anti-mouse CD8a | 53-6.7 | BioLegend | 100708 |
| Alexa Fluor® 488 anti-mouse CD80 | 16-10A1 | BioLegend | 104716 |
| Brilliant Violet 605™ anti-mouse CD86 | GL-1 | BioLegend | 105037 |
| BD Horizon™ PE-CF594 Hamster Anti-Mouse T-Cell Receptor | GL3 | BD Biosciences | 563532 |
| BD Horizon™ BUV496 Rat Anti-Mouse CD4 | GK1.5 | BD Biosciences | 612952 |
| BD Horizon™ BUV395 Hamster Anti-Mouse TCR Chain | H57-597 | BD Biosciences | 569248 |
| BD Horizon™ PE-CF594 Rat Anti-Mouse Siglec-F | E50-2440 | BD Biosciences | 562757 |
| BD Horizon™ BUV737 Rat Anti-CD11b | M1/70 | BD Biosciences | 612800 |
| BD Horizon™ BUV805 Rat Anti-Mouse CD45 | 30-F11 | BD Biosciences | 568336 |

**Table S11.** List of antibody markers for flow cytometry.
